# ARHGEF3 regulates skeletal muscle regeneration and strength through autophagy

**DOI:** 10.1101/2020.02.28.970756

**Authors:** Jae-Sung You, Nilmani Singh, Adriana Reyes-Ordonez, Nidhi Khanna, Zehua Bao, Huimin Zhao, Jie Chen

## Abstract

Skeletal muscle regeneration is essential for restoring muscle function upon injury and for the maintenance of muscle health with aging. ARHGEF3, a Rho-specific GEF, negatively regulates myoblast differentiation via mammalian target of rapamycin complex 2 (mTORC2)-Akt signaling in a GEF-independent manner *in vitro*. Here, we investigated ARHGEF3’s role in skeletal muscle regeneration by creating ARHGEF3 KO mice. These mice exhibited no discernible phenotype under normal conditions. Upon injury, however, ARHGEF3 deficiency enhanced the mass, fiber size and function of regenerating muscles in both young and aged mice. Surprisingly, these effects were not mediated by mTORC2-Akt signaling, but by the GEF activity of ARHGEF3. Furthermore, ARHGEF3 KO promoted muscle regeneration through activation of autophagy, a process that is also critical for maintaining muscle strength. Accordingly, in old mice, ARHGEF3 depletion prevented muscle weakness by restoring autophagy flux. Collectively, our findings identify an unexpected link between ARHGEF3 and autophagy-related muscle pathophysiology.

## Introduction

Skeletal muscle possesses robust regeneration capacity that is essential for restoration of intact muscle function upon injury. Successful muscle regeneration requires a highly coordinated myogenesis process consisted of muscle stem cell activation and proliferation, cell-cycle exit, and fusion of mono-nucleated myoblasts (Le Grand and Rudnicki, 2007). Although many signaling molecules that modulate these processes have been identified *in vitro*, their role in muscle regeneration *in vivo* is still largely unexplored. Nevertheless, some intracellular mechanisms critical for muscle regeneration have recently been identified. For example, autophagy, a key homeostatic process necessary for degradation of unwanted cellular components, plays a crucial role in successful muscle regeneration after injury (Call et al., 2017; Garcia-Prat et al., 2016; Paolini et al., 2018). Again, however, molecules that regulate autophagy during regeneration are poorly understood.

More than 10 years ago, many research groups had been interested in and investigated the role of RhoA/Rho-associated kinase (ROCK) signaling in myogenic differentiation, and their studies came to the conclusion that RhoA/ROCK regulation of myogenesis in cells is cell stage-specific: it is necessary for myoblast proliferation and maintenance of myogenic capacity, but it suppresses differentiation once cells exit the cell cycle and enter the post-proliferative phase (Charrasse et al., 2006; Iwasaki et al., 2008; Lim et al., 2007; Takano et al., 1998; Wei et al., 1998). Although these studies suggested the potential involvement of a guanine nucleotide exchange factor for Rho GTPases (RhoGEF) in skeletal muscle regeneration, there has been no report of a role of any endogenous RhoGEF (among ∼70 in the human genome (Rossman et al., 2005)) in muscle regeneration. Perhaps targeting RhoA signaling with such complex and paradoxical effects is presumed to be less clinically feasible and effective.

Since then, Khanna et al. had examined ARHGEF3 (also called XPLN), which selectively activates RhoA and RhoB (Arthur et al., 2002), and found that this RhoGEF negatively regulates myoblast differentiation through inhibition of the mammalian target of rapamycin (mTOR) complex 2 (mTORC2) activation of Akt (Khanna et al., 2013), one of the best established myogenic factors (Jiang et al., 1999). Interestingly, in this study, ARHGEF3 exerted these functions independently of its GEF activity, thus raising the possibility that ARHGEF3 could become a mechanistically feasible target for regulating skeletal muscle regeneration regardless of its role in RhoA signaling. In the current study, we investigated this possibility by creating ARHGEF3 knockout (KO) mice. These mice showed no discernible phenotype during development. However, depletion of ARHGEF3 not only promoted injury-induced muscle regeneration but also prevented age-related regenerative defects. Unexpectedly, these effects of ARHGEF3 KO did not require activation of mTORC2-Akt signaling, but instead they occurred through a mechanism involving the GEF activity of ARHGEF3 and autophagy.

## Results

### ARHGEF3 KO promotes skeletal muscle regeneration after injury

*Arhgef3* gene-edited mice were created by TALEN-mediated targeting of exon 3, the earliest region shared by all four established *Arhgef3* transcript variants, in the germ line on a C57BL/6N background. A mouse line containing a 17-base pair deletion in exon 3 (thus causing open reading frame shift) was selected for further characterization. Mice were bred to yield homozygous mutant offspring, identified by genotyping with three independent PCR-based methods (Figure 1A). Muscles from the mutant mice expressed a drastically reduced level of *Arhgef3* transcripts (Figure 1B, left panel) most likely due to nonsense-mediated RNA decay. Furthermore, sequencing of the remaining transcripts confirmed the frame-shift deletion of exon 3 and introduction of premature stop codons (Figure 1B, right panel). Only the first 58aa of ARHGEF3 would be translated from this mutant mRNA, which is unlikely to result in a functional or even stable protein. From here on, we refer to the homozygous mutant mice as ARHGEF3 KO. These mice did not exhibit any discernible phenotypic abnormality during development (Figure 1C).

**Figure 1.**
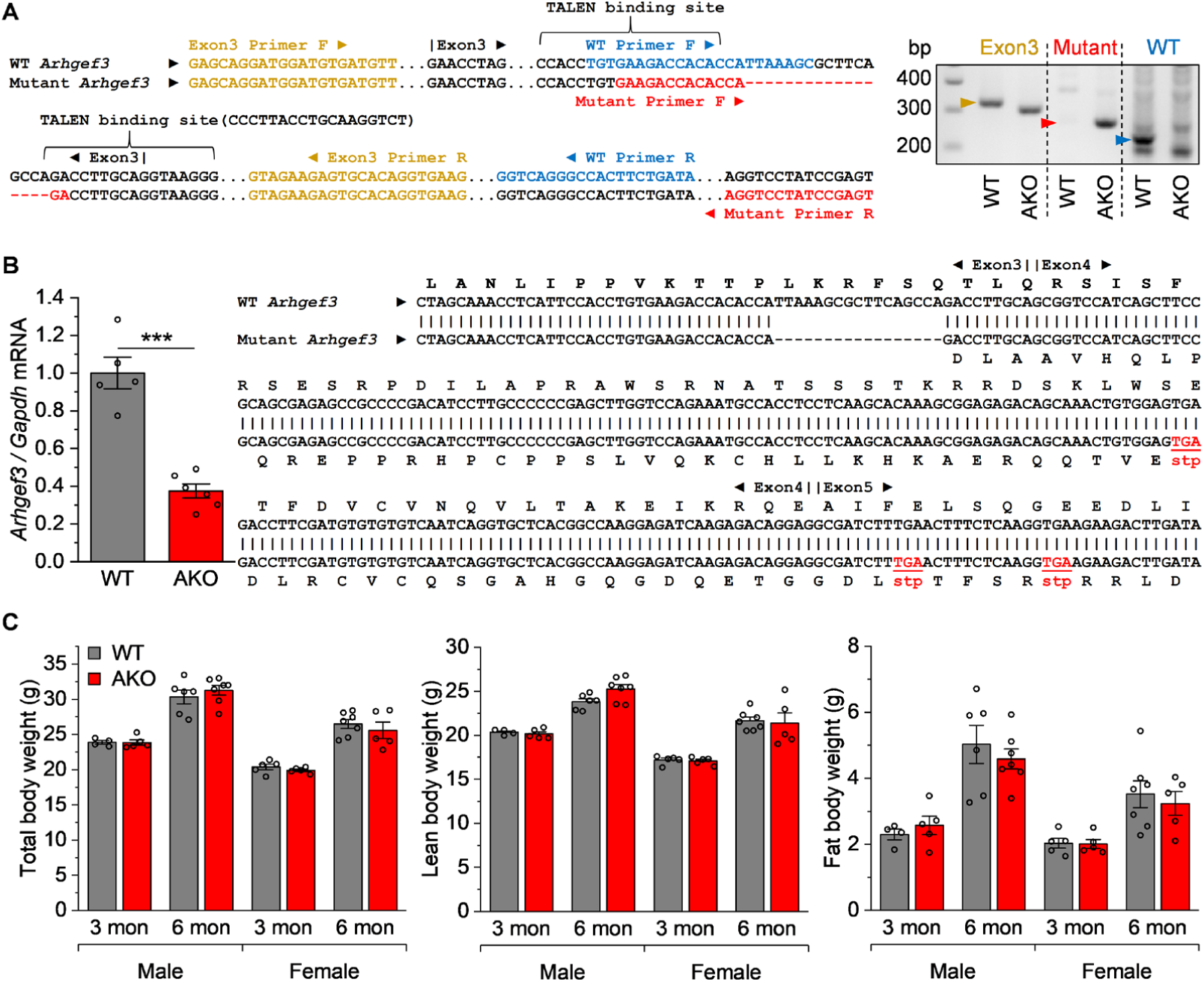
Generation of ARHGEF3 KO mice. (**A**) Genomic DNA extracted from TA muscles of WT and ARHGEF3 KO (AKO) mice was PCR-amplified with three different primer sets (left panel) and the amplicons were detected by agarose gel electrophoresis (right panel). (**B**) cDNA from mRNA extracted from TA muscles of WT and AKO mice was subjected to quantitative PCR analyses (left panel, *n* = 5-6) or sequenced (right panel). (**C**) Body compositions were analyzed in 3- and 6-month (mon)-old WT and AKO mice (*n* = 4-7). Data are presented as mean ± SEM. \*\*\**P* < 0.001 by 2-tailed unpaired *t* test.

To address the role of ARHGEF3 in skeletal muscle regeneration, tibialis anteria (TA) muscles from WT and ARHGEF3 KO mice were injected with BaCl_2_, which induces muscle fiber necrosis and subsequent regeneration. Upon injury, the ARHGEF3 protein was transiently elevated in WT, but not in ARHGEF3 KO, muscles (Figure 2A) allowing its detection by Western blotting and further confirmation of ARHGEF3 KO. Detection of ARHGEF3 by Western has been a long-standing challenge. We tested nearly all commercially available antibodies and none was validated in cells by ARHGEF3 knockdown. Our custom-made antibody used here was the only one validated (Khanna et al., 2013) although its avidity is not high.

**Figure 2.**
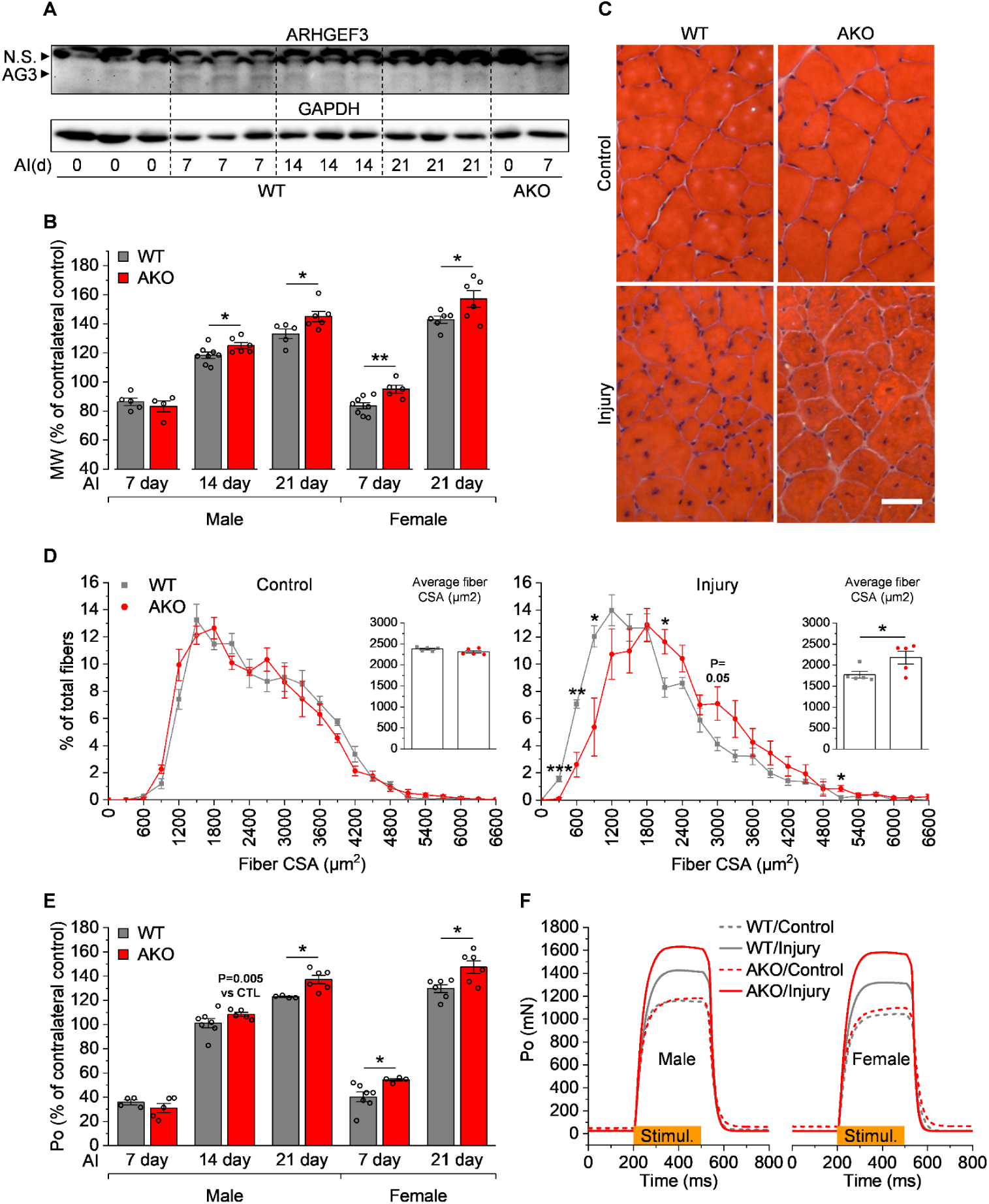
ARHGEF3 KO promotes skeletal muscle regeneration after injury. TA muscles from 3-month-old WT and ARHGEF3 KO (AKO) mice were injected with BaCl_2_ (injury) or saline (uninjured control, 0 day). (**A**) Protein expression of ARHGEF3 (AG3) and GAPDH was analyzed by Western blotting in female muscles collected 7, 14, and 21 days (d) after injury (AI) (*n* = 3). N.S., non-specific. (**B**) Muscle weight (MW) was measured 7, 14, and 21 days after AI and presented as % of contralateral uninjured control (*n* = 4-8). (**C**) Representative H&E images of cross sections of injured and uninjured muscles collected 21 days AI from male mice. Scale bar: 50 µm. (**D**) Cross-sectional area (CSA) of myofibers was measured from the H&E images and presented on histograms with inserts showing averaged CSA (*n* = 5). (**E**) Maximal isometric tetanic force (Po) was measured 7, 14, and 21 days AI and presented as % of contralateral uninjured control (CTL) (*n* = 4-7). (**F**) Representative trace of maximal isometric tetanic force during stimulation (300ms) at 21 days AI. Data are presented as mean ± SEM. \**P* < 0.05, \*\**P* < 0.01 by 2-tailed unpaired *t* test unless otherwise indicated.

Over the course of regeneration, muscle mass increased steadily in WT mice as expected, and this increase was further enhanced in both male and female ARHGEF3 KO mice (Figure 2B), suggesting that KO of ARHGEF3 promoted muscle regeneration. To further assess regeneration, we analyzed cross-section area of the newly regenerating myofibers identified by their central nuclei (Figure 2C), and found that KO of ARHGEF3 led to a shift of regenerating myofibers to larger sizes, resulting in a significant increase in the average size (Figure 2D).

Next, we asked whether those morphological effects of ARHGEF3 KO were functionally relevant. To address that, we measured muscle force *in situ* over the course of regeneration. Similar to muscle mass changes, total muscle force progressively increased after injury and ARHGEF3 KO further enhanced the force in both male and female mice with more prominent effects after 7 days post-injury (Figures 2E and 2F). These results clearly demonstrate that depletion of the ARHGEF3 protein promotes skeletal muscle regeneration, in particular the post-proliferative phase of regeneration (Tidball and Villalta, 2010), *in vivo*. Since we did not find any fundamental gender difference in the injury phenotype between WT and ARHGEF3 KO mice, we pursued next experiments with one gender whichever available.

### ARHGEF3 regulation of muscle regeneration is independent of mTORC2-Akt signaling

Next, we wanted to determine the mechanism underlying the effects of ARHGEF3 KO on muscle regeneration. Previously, it had been shown that knockdown of ARHGEF3 increases myogenic mTORC2-Akt signaling in C2C12 myoblasts (Khanna et al., 2013). To investigate the involvement of mTORC2-Akt signaling, we examined phosphorylation status of mTORC2 substrates including Ser473-Akt, Ser657-PKCα (Ikenoue et al., 2008), and SGK1 (Garcia-Martinez and Alessi, 2008), in BaCl_2_-injected regenerating muscles. As shown in Figures 3A and S1, ARHGEF3 KO increased phosphorylation of Ser473-Akt, but not Thr308-Akt, Ser657-PKCα, or Thr346-NDRG1 (a substrate of SGK1). These observations are consistent with the previous *in vitro* finding that ARHGEF3 specifically inhibits mTORC2 phosphorylation of Akt (Khanna et al., 2013).

**Figure 3.**
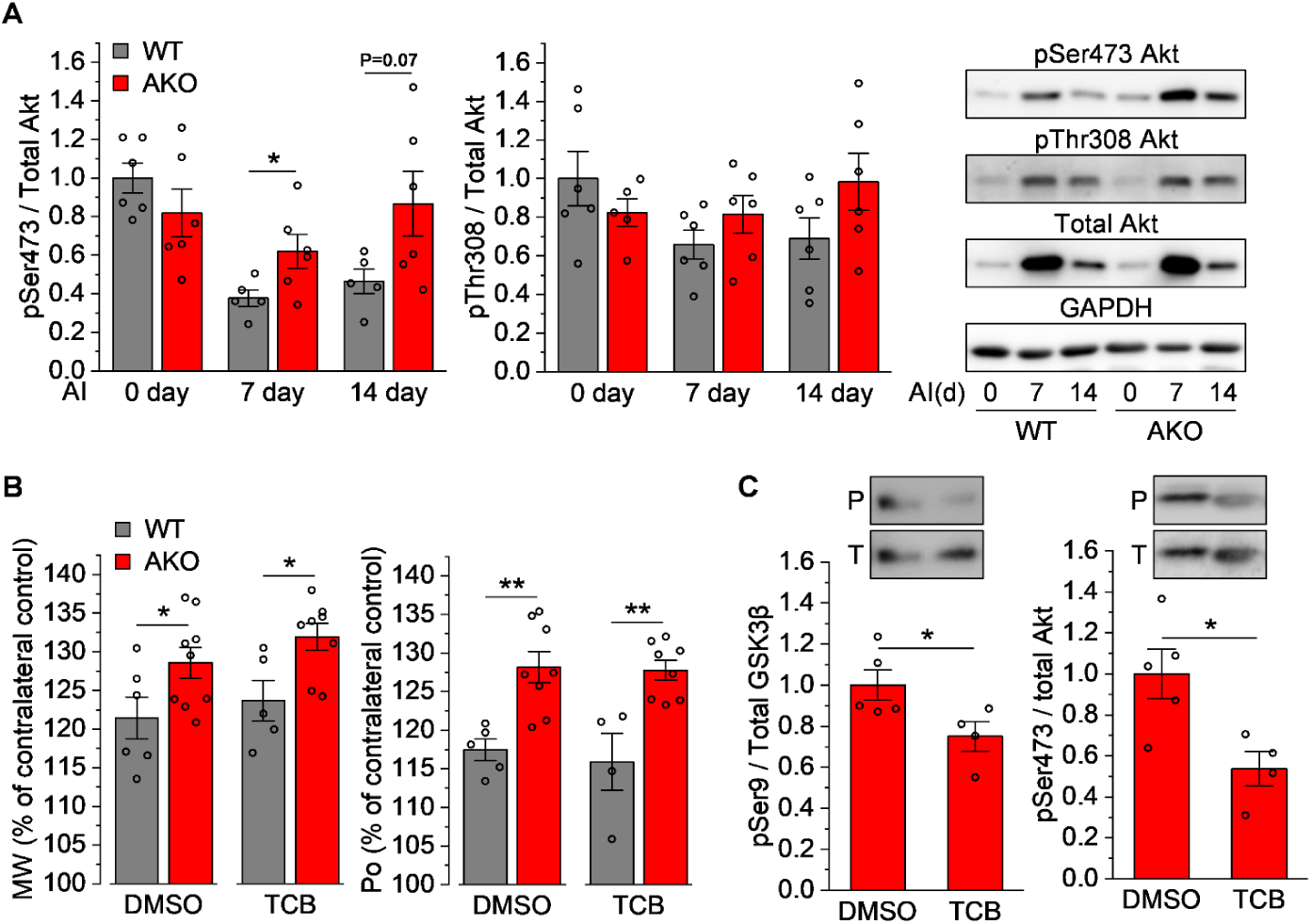
ARHGEF3 regulation of muscle regeneration is independent of mTORC2-Akt signaling. TA muscles from 3-month-old WT and ARHGEF3 KO (AKO) male mice were injected with BaCl_2_ (injury) or saline (uninjured control, 0 day). (**A**) The muscles were collected 7 and 14 days after injury (AI) and analyzed by Western blotting for phosphorylated (p)/total Akt ratio and GAPDH protein expression (*n* = 5-6). (**B**-**C**) Mice were treated with vehicle (DMSO) or triciribine (TCB) upon injury. Muscle weight (MW) and maximal isometric tetanic force (Po) were measured 21 days AI and presented as % of contralateral uninjured control (**B**, *n* = 4-9). Phosphorylated (P)/total (T) protein ratio for Akt and GSK3β was determined by Western blotting in injured AKO muscles collected 21 days AI (**C**, *n* = 4-5). Data are presented as mean ± SEM. \**P* < 0.05, \*\**P* < 0.01 by 2-tailed unpaired *t* test (A, C) or 2-way ANOVA (B). See also Figure S1.

To directly test whether the effects of ARHGEF3 KO on muscle regeneration is mediated by mTORC2-Akt signaling, we treated mice with the Akt inhibitor triciribine, which has been shown to be highly effective in blocking Akt-dependent increase in skeletal muscle regeneration (Waldemer-Streyer and Chen, 2015). As expected, muscles from ARHGEF3 KO mice treated with control vehicle displayed increased muscle mass and force compared to that from WT mice after BaCl_2_ injection (Figure 3B). Surprisingly however, triciribine did not prevent any of these effects of ARHGEF3 KO (Figure 3B) despite substantial inhibition of phosphorylation of Akt and its substrate GSK3β (Figure 3C). Hence, we conclude that ARHGEF3 KO promotes muscle regeneration through a mechanism independent of the mTORC2-Akt signaling pathway.

### The GEF activity of ARHGEF3 is critical for the regulation of muscle regeneration

The observation that ARHGEF3 KO exerted more prominent effects on regeneration after the proliferative stage (0-7days) (Figures 1B and 1E) is consistent with the cell stage-specific role of RhoA/ROCK signaling in myogenesis *in vitro* (Iwasaki et al., 2008). Hence, we considered a potential involvement of the GEF activity of ARHGEF3 in the regulation of muscle regeneration. Indeed, we found that RhoA-specific GEF activity was substantially reduced by ARHGEF3 KO in regenerating muscles (Figure 4A). To further probe a functional relevance of the GEF activity, we introduced recombinant ARHGEF3, WT or a GEF-inactive mutant (L269E) (Khanna et al., 2013) by *in vivo* transfection into the regenerating TA muscles at 10 days post-injury, a time point after which the muscles undergo the post-proliferative phase of regeneration (Figure 4B). Furthermore, at this time point, F4/80-positive infiltrating macrophages, which may complicate the interpretation of our experiments, remarkably subsided (Figure 4B). As shown in Figure 4C, the ARHGEF3 KO muscles transfected with L269E-ARHGEF3 displayed lower RhoA GEF activity than that with WT-ARHGEF3, and the difference was very similar to that between intact WT and ARHGEF KO regenerating muscles (Figure 4A). However, unlike the WT and ARHGEF3 KO muscles, this difference in RhoA GEF activity did not affect Ser473-Akt phosphorylation (Figure 4C), which is consistent with the GEF-independent role of ARHGEF3 on Akt phosphorylation (Khanna et al., 2013). Strikingly, the absence of ARHGEF3 GEF activity led to better muscle regeneration as indicated by bigger mass and increased force production of the regenerating muscles (Figure 4D). Collectively, these results indicate that the GEF activity of ARHGEF3 is critical for the regulation of muscle regeneration.

**Figure 4.**
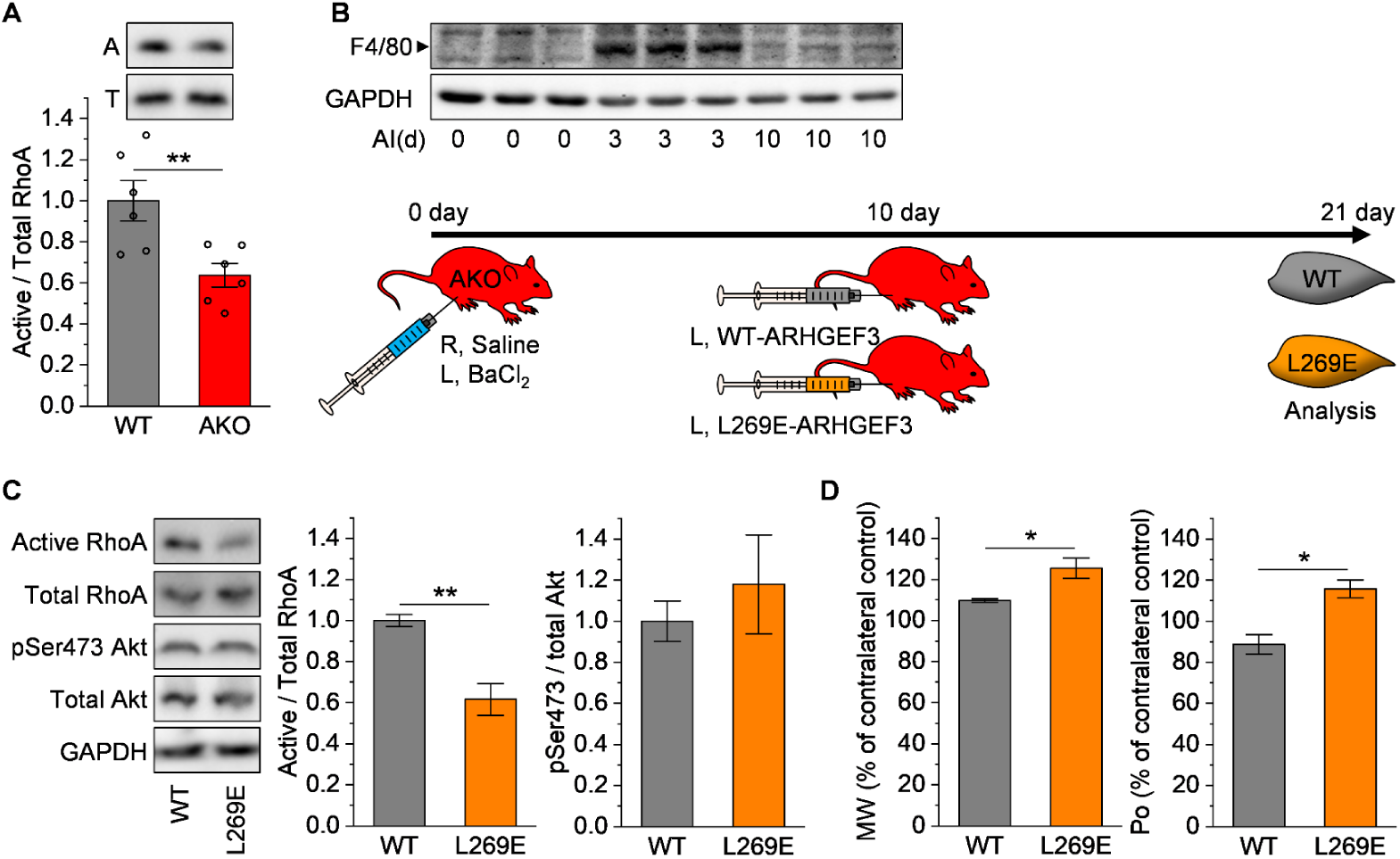
The GEF activity of ARHGEF3 is critical for the regulation of muscle regeneration. (**A**) TA muscles from 3-month-old WT and ARHGEF3 KO (AKO) male mice were injected with BaCl_2_, collected 14 days after injury, and analyzed by Western blotting for active (A)/total (T) RhoA ratio (*n* = 5-6). (**B**) TA muscles from ARHGEF3 KO (AKO) male mice were injected with BaCl_2_ (injury) or saline (uninjured control, 0 day), and either collected 3 and 10 days (d) after injury (AI) for analysis of F4/80 and GAPDH protein expression by Western blotting (upper panel, *n* = 3) or transfected with plasmids encoding WT- or L269E-ARHGEF3 10 days AI and allowed to recovery until 21 days AI (lower panel). (**C**) Active/total RhoA ratio, phosphorylated (p)/total Akt ratio, and GAPDH protein expression were determined by Western blotting in the transfected injured muscles (*n* = 3). (**D**) Muscle weight (MW) and maximal isometric tetanic force (Po) were measured in the transfected injured muscles and presented as % of contralateral uninjured control (*n* = 3). Data are presented as mean ± SEM. \**P* < 0.05, \*\**P* < 0.01 by 2-tailed unpaired *t* test.

### ARHGEF3 KO promotes muscle regeneration through autophagy and prevents age-related regenerative defects

Recently, a potential link between RhoA signaling and autophagy has been reported in mouse cardiomyocytes (Shi et al., 2018; Shi et al., 2019): deletion of ROCK1, a major downstream effector of RhoA, restored autophagy flux that was impaired by doxorubicin, a broad-spectrum anti-cancer drug (Shi et al., 2018), ROCK1/ROCK2 double deletion promoted basal autophagy and reduced cardiac fibrosis during aging (Shi et al., 2019). In light of the pivotal role of GEF activity in mediating ARHGEF3 KO effects and this reported link between RhoA signaling and autophagy, we hypothesized that ARHGEF3 KO might promote autophagy during muscle regeneration. As such, we monitored autophagy flux by using autophagy marker LC3-II, a lipidated form of LC3. LC3-II was elevated in muscles injected with BaCl_2_ and this increase was further augmented by the treatment of the autophagy inhibitor chloroquine; the effect of chloroquine was similar in injured and uninjured control muscles (Figures 5A and S2A). Because the level of LC3-II accumulation by autophagy blockage reflects the level of autophagy flux, these results indicate that autophagy flux was not altered during regeneration. We then compared WT and ARHGEF3 KO muscles and found that the level of LC3-II, but not GAPDH, was significantly reduced by ARHGEF3 KO in injured muscles (Figures 5B and S2B, left panels). To determine whether this reduction was due to increased autophagic degradation of LC3-II or decreased conversion of LC3-I into LC3-II, we also compared injured muscles from WT and ARHGEF3 KO after chloroquine treatment and found no difference in the level of LC3-II (Figures 5B and S2B, right panels). Hence, these results indicate that ARHGEF3 KO increased autophagy flux and thus the autophagic degradation of LC3-II after injury. In support of this notion, ARHGEF3 KO also reduced the protein level of another autophagy-selective substrate p62 in injured muscles without affecting its mRNA levels (i.e., increased autophagic degradation of p62) (Figures 5C and 5D).

**Figure 5.**
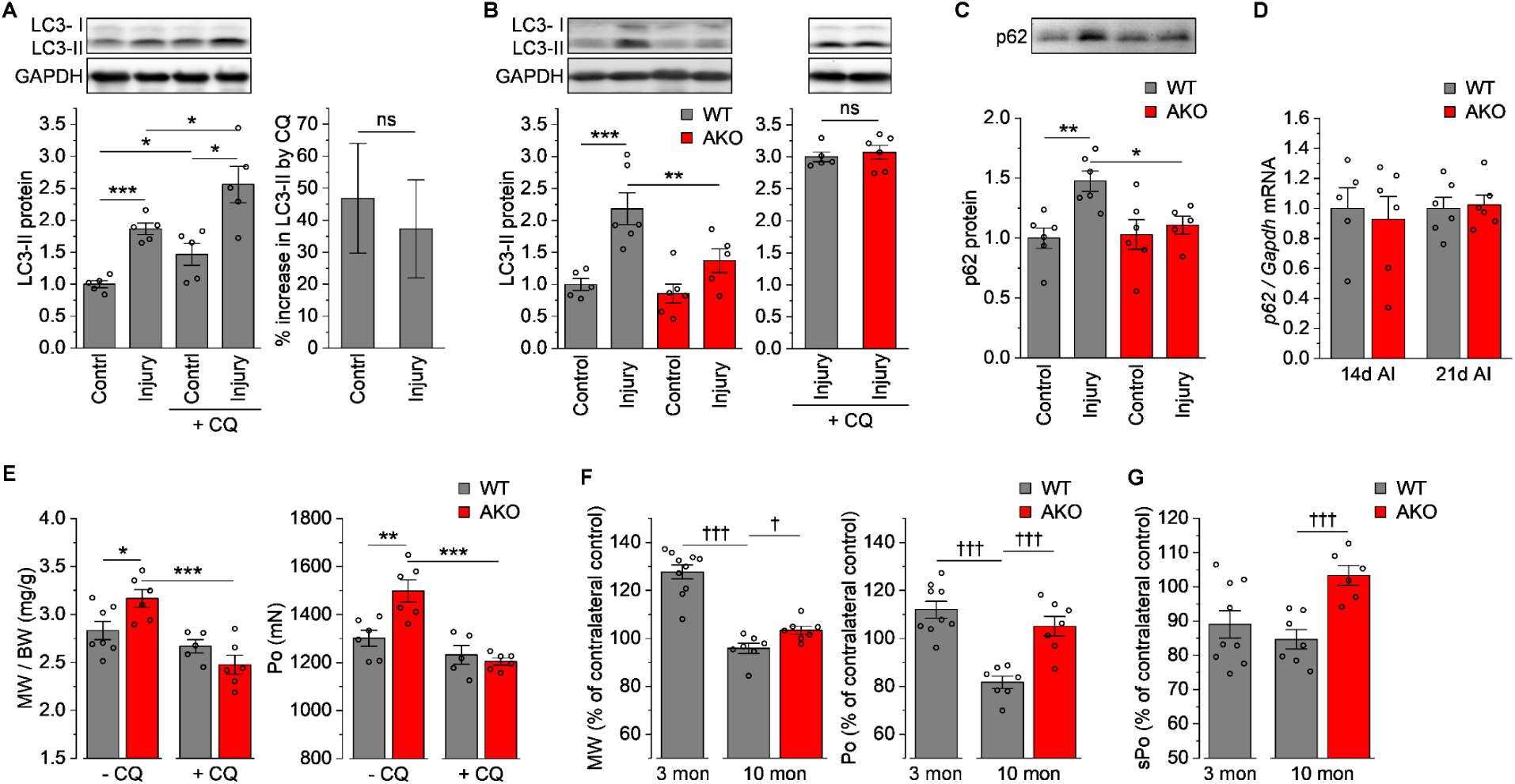
ARHGEF3 KO promotes muscle regeneration through autophagy and prevents age-related regenerative defects. TA muscles from WT and/or ARHGEF3 KO (AKO) female mice were injected with BaCl_2_ (injury) or saline (uninjured control) and allowed to recover with or without chloroquine (CQ) treatment. (**A**) Protein expression of LC3-II and GAPDH was analyzed by Western blotting in 3-month-old WT muscles collected 21 days after injury (*n* = 5). (**B**-**C**) Protein expression of LC3-II, GAPDH (**B**), and p62 (**C**) was analyzed by Western blotting in 3-month-old WT and AKO muscles collected 21 days after injury (*n* = 5-6). (**D**) mRNA expression of p62 was analyzed by qPCR in 3-month-old WT and AKO muscles collected 14 and 21 days (d) after injury (AI) (*n* = 5-6). (**E**) Muscle weight (MW)/body weight (BW) ratio and maximal isometric tetanic force (Po) were measured in 3-month-old WT and AKO injured muscles 21 days after injury (*n* = 5-7). (**F**-**G**) Muscle weight (MW), maximal isometric tetanic force (Po) (**F**), and specific maximal isometric tetanic force (sPo) (**G**) were measured in 3- and/or 10-month-old WT and AKO muscles 14 days after injury and presented as % of contralateral uninjured control (*n* = 6-10). Data are presented as mean ± SEM. \**P* < 0.05, \*\**P* < 0.01, \*\*\**P* < 0.001 by 2-way ANOVA. ^†^*P* < 0.05, ^†††^*P* < 0.001 by 2-tailed unpaired *t* test. See also Figure S2.

We then asked if this increase in autophagy flux by ARHGEF3 KO was responsible for the enhanced muscle regeneration in ARHGEF3 KO muscles. As shown in Figure 5E, inhibition of autophagy by chloroquine abolished the ARHGEF3 KO-induced increases in muscle mass and force after injury. Therefore, it is highly likely that ARHGEF3 KO promotes muscle regeneration by enhancing autophagy after injury.

Skeletal muscle regenerative capacity declines with aging owing to a suppression of autophagy in multiple cell types (Lee et al., 2019). Hence, we envisioned that ARHGEF3 KO with autophagy-promoting effect would counteract the age-related decline in muscle regenerative capacity. As shown in Figure 5F, 10-month-old mice clearly exhibited retarded recovery in muscle mass and total muscle force after BaCl_2_-induced injury when compared to 3-month-old mice. Importantly, these regenerative defects were significantly ameliorated in ARHGEF3 KO mice (Figure 5F). We also found that, unlike in young animals (Figure 2B and 2E), the rescue effects were particularly prominent in total muscle force and that this was due to a robust recovery of injured muscle quality (specific muscle force) by ARHGEF3 KO. Together, these results suggest that ARHGEF3 may be a potential target for preserving regenerative capacity during aging.

### ARHGEF3 KO prevents age-related muscle weakness by restoring autophagy flux

The effect of ARHGEF3 KO on regenerating muscle quality in aged mice is particularly interesting because, even in the absence of injury, skeletal muscle undergoes a loss of muscle strength with aging (dynapenia) and this occurs to a much greater degree than age-related loss of muscle mass (sarcopenia) (Goodpaster et al., 2006; Metter et al., 1999; Russ et al., 2012). An emerging body of evidence also strongly suggests that this age-related loss of specific muscle strength, or muscle quality, is attributable to a decline in basal autophagy and subsequent defects in force transmission apparatus including neuromuscular junction (Bujak et al., 2015; Carnio et al., 2014; Demontis and Perrimon, 2010; Sebastian et al., 2016). Hence, it is plausible to consider a link between ARHGEF3, autophagy, and muscle quality in aging.

To further establish a link between autophagy and muscle quality, we performed an experiment where young mice received a single dose of chloroquine for time course examination of muscle quality. As shown in Figure 6A, chloroquine rapidly induced a transient loss of specific muscle force, and its temporal pattern was in a close inverse relationship with the level of LC3-II accumulation, as well as the twitch/tetanic force ratio representing innervation ratio (Celichowski and Grottel, 1993). Of note, innervation ratio increases with aging (Brooks and Faulkner, 1988; Fling et al., 2009; Kung et al., 2014).

**Figure 6.**
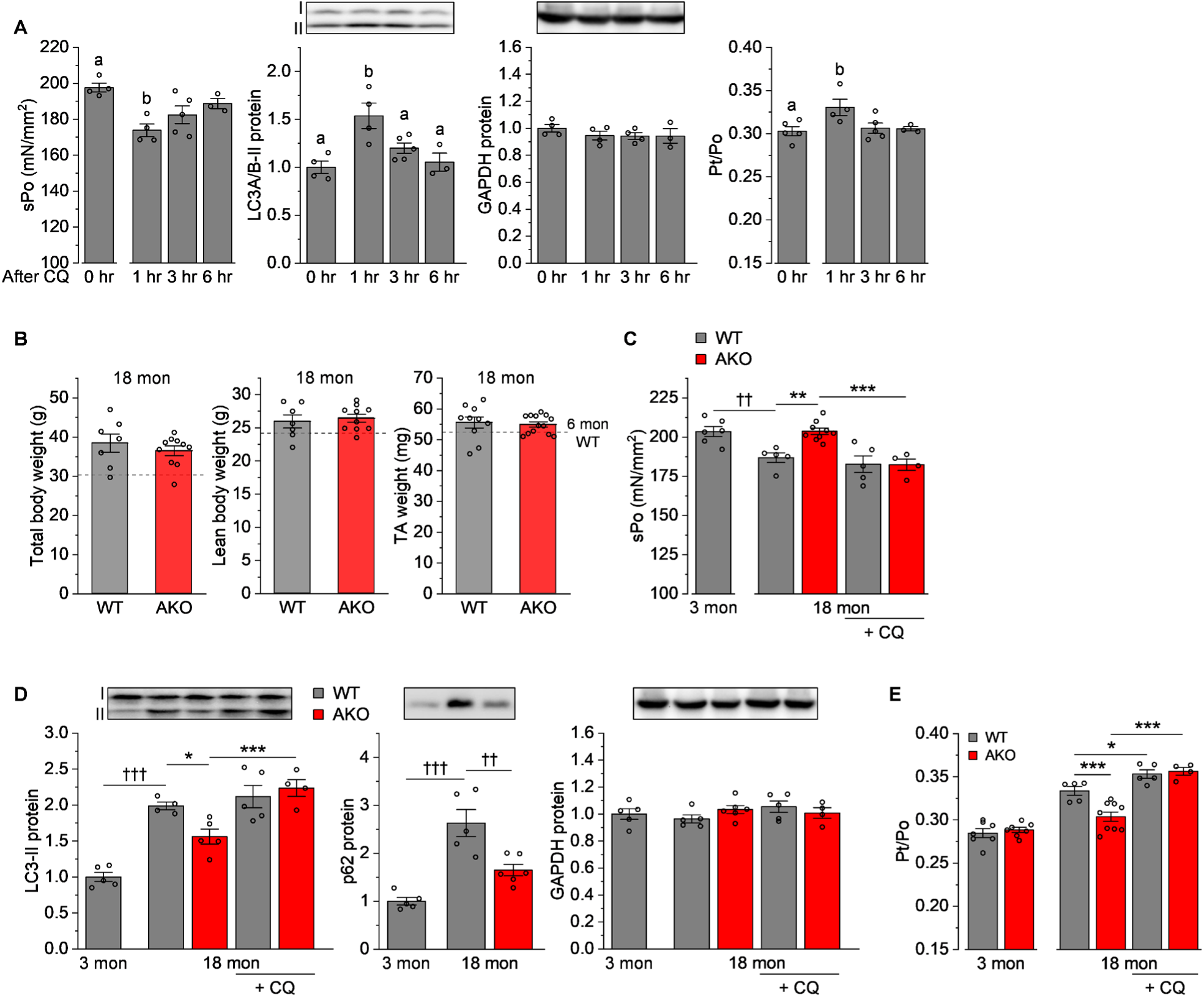
ARHGEF3 KO prevents age-related muscle weakness by restoring autophagy. (**A**) Three-month-old mice were injected with vehicle (0 hr) or chloroquine (CQ) and, at indicated time points, TA muscles were analyzed for specific maximal isometric tetanic force (sPo) and collected for analysis of LC3-II and GAPDH protein expression by Western blotting (*n* = 3-5). The muscles were also analyzed for maximal twitch force (Pt)/maximal tatanic force (Po) ratio (*n* = 3-5). (**B**) Eighteen-month-old WT and ARHGEF3 KO (AKO) male mice were analyzed for total body weight, lean body weight, and TA muscle weight (*n* = 7-13). Dashed lines represent mean values from 6-month (mon)-old WT male mice (*n* = 6-7). (**C**-**E**) Three- and/or 18-month-old WT and AKO male mice were injected with vehicle or CQ and, at 1 hour after the injection, TA muscles were analyzed for sPo (**C**, *n* = 4-9) and collected for analysis of LC3-II, p62, and GAPDH protein expression by Western blotting (**D**, *n* = 4-6). The muscles were also analyzed for Pt/Po ratio (**E**, *n* = 4-9). Data are presented as mean ± SEM. ^a,b^*P* < 0.05 from each other by 1-way ANOVA, ^††^*P* < 0.01, ^†††^*P* < 0.001 by 2-tailed unpaired *t* test, \**P* < 0.05, \*\**P* < 0.01, \*\*\**P* < 0.001 by 2-way ANOVA.

Then, we went on to address whether ARHGEF3 KO could preserve muscle quality in old mice by modulating autophagy. When compared to fully grown 6-month-old adult mice, 18-month-old WT mice did not show any evidence of age-related muscle loss as previously reported (Hamrick et al., 2006), and nor did ARHGEF3 KO mice (Figure 6B). However, specific muscle force was significantly diminished by aging in WT mice and this decline in muscle quality was completely prevented in ARHGEF3 KO mice (Figure 6C). We also found that ARHGEF3 KO reduced aging-induced accumulation of LC3-II and p62 (Figure 6D). Unlike in young mice, a single dose of chloroquine did not further decrease specific muscle force or increase LC3-II level in old WT mice, confirming that these mice were already suffering from severe autophagy inhibition (Figures 6C and 6D). However, in old ARHGEF3 KO mice, chloroquine both decreased specific muscle force and increased the level of LC3-II, leading to a complete reversal of the rescue effects of ARHGEF3 KO on aging-induced deterioration in muscle quality and basal autophagy (Figures 6C and 6D). The same pattern of changes was also observed in twitch/tetanic force ratio (Figure 6E). Overall, these results demonstrate that depletion of ARHGEF3 allows maintenance of basal autophagy flux and thereby preserves muscle quality in aging muscle.

## Discussion

Skeletal muscle’s ability to regenerate has an immediate clinical relevance in pathological and non-pathological conditions such as muscular dystrophies, post-injury recovery, and aging. In this study, by using TALEN-mediated gene editing, we demonstrated that depletion of ARHGEF3, particularly its GEF activity, promoted functional muscle regeneration after injury in both young and aged mice. We also found that depletion of ARHGEF3 rescued specific muscle strength in old mice, and that all these effects were driven by autophagy activation. Hence, our findings uncover ARHGEF3 as a novel regulator of muscle regeneration and muscle quality, and highlight ARHGEF3 as a potential therapeutic target for impaired muscle regeneration and function as well as other clinical issues that result from autophagy defects (Levine and Kroemer, 2008).

Based on the current knowledge derived from *in vitro* studies, endogenous ARHGEF3 exerts two molecular functions, one as a RhoGEF toward RhoA/B and the other as an inhibitor of mTORC2-Akt signaling (Arthur et al., 2002; Khanna et al., 2013). In agreement with those activities, we have found that ARHGEF3 KO resulted in reduced RhoA GEF activity and increased mTORC2-Akt signaling in regenerating mouse skeletal muscle. Hence, we examined involvement of both mechanisms and found that RhoGEF activity, rather than the well-established myogenic mTORC2-Akt signaling, mediates the regeneration-promoting effects of ARHGEF3 KO. Furthermore, our findings implicate autophagy in the pathway through which ARHGEF3/Rho signaling regulates muscle regeneration. This mechanism is consistent with recent reports that ROCK deletion enhanced autophagy in cardiomyocytes (Shi et al., 2018; Shi et al., 2019). To further understand ARHGEF3 regulation of muscle regeneration, future investigations aimed at defining how ARHGEF3/Rho signaling regulates autophagy during regeneration are needed.

Autophagy is also suppressed by Akt via its multiple downstream effectors including mTORC1, FoxOs, and Beclin1 (Mammucari et al., 2008; Wang et al., 2012). Because ARHGEF3 both activates RhoA upstream of ROCK and inhibits Akt, it is reasonable to propose a model where depletion of AGHGEF3 enhances autophagy by reducing the inhibitory RhoA/ROCK signaling while simultaneously suppressing autophagy with Akt activation. Although these two opposing effects of autophagy are expected to counteract each other, our observation of autophagy induction in regenerating muscle with AGHGEF3 depletion indicates that ARHGEF3 acts more predominantly on the RhoA/ROCK-autophagy pathway than the Akt-autophagy pathway during regeneration. However, we also speculate that, once this Akt-mediated autophagy suppression mode is released by an Akt inhibitor such as triciribine, autophagy would be further activated in ARHGEF3 KO muscles, offsetting any negative effect the Akt inhibitor may have on myogenesis.

In this study, depletion of ARHGEF3 in mice did not result in any phenotype under normal conditions, but it exerted beneficial effects in the contexts of injury and aging through activation of autophagy. These results suggest that the activity of ARHGEF3 toward autophagy inhibition is switched on only under stressful conditions such as muscle injury or aging. This mode of action may make ARHGEF3 a particularly suitable therapeutic target to allow induction of autophagy specifically in afflicted tissues without the side effects of unnecessary boosting of autophagy in non-targeted or healthy cells/tissues that could disrupt cellular homeostasis (Thorburn, 2018). An interesting question is how ARHGEF3 activity is up-regulated under those specific conditions. It appears that skeletal muscle expresses a relatively low level of ARHGEF3 transcript when compared to other tissues and other Rhos GEFs (Cario-Toumaniantz et al., 2012). Hence, one immediate speculation is that the expression of ARHGEF3 is induced under the specific conditions. Indeed, the ARHGEF3 protein was up-regulated upon injury. However, this mechanism is less likely to be responsible for the regulation of muscle regeneration because the effect of ARHGEF3 KO on regeneration was more prominent at a time when the ARHGEF3 protein level had returned to its basal state. An alternative possibility is the regulation of ARHGEF3 activity by subcellular localization. A recent study has found that ARHGEF3 is normally enriched in the nucleus and it translocates into the cytoplasm upon inhibition of histone deacetylases, leading to activation of RhoA and actin cytoskeletal reorganization (D’Amato et al., 2015). Therefore, the RhoGEF activity of ARHGEF3 may be very low under normal conditions due to its nuclear sequestration, and it is switched on when specific conditions induce ARHGEF3 export from the nucleus. Post-translational modifications such as phosphorylation may also regulate RhoGEF activity of ARHGEF3, as reported for other RhoGEFs (Patel and Karginov, 2014). It will be of great importance for future studies to probe these mechanistic questions in order to elaborate therapeutic strategies targeting ARHGEF3.

## Acknowledgements

We thank Fuming Pan and the Transgenic Mouse Facility at the University of Illinois at Urbana-Champaign for pronuclear injection of TALEN mRNA and creation of founder transgenic mice. This work was supported by a grant from the National Institutes of Health to JC (R01AR048914).

## Author Contributions

J.-S.Y. and J.C. designed the study; Z.B. and H.Z. designed and created the TALENs; N.S., A.R., N.K., and J.-S.Y. generated mouse lines; J.-S.Y. acquired and analyzed data; J.-S.Y. and J.C. wrote the manuscript; N.S., N.K. and H.Z. edited the manuscript.

## Methods

### Generation of ARHGEF3 KO

We constructed five pairs of pCS2+TALEN plasmids targeting different regions of exon 3 of the mouse *Arhgef3* gene. Each pair of the plasmids was transfected in C2C12 myoblasts, and genomic DNA was extracted after 3 days of transfection. The genomic region encompassing the TALEN binding sites was PCR-amplified and screened for successful mutations by using the SURVEYOR Mutation Detection Kit (Transgenomic). A TALEN pair selected for *in vivo Arhgef3* editing (see Figure 1A for TALEN binding sites) was linearized with ApaI and transcribed into mRNA using the mMESSAGE mMACHINE SP6 transcription Kit (Invitrogen). The RNA was purified with the MEGAclear Transcription Clean-Up Kit (Invitrogen) and resuspended in T10E0.1 injection buffer. The RNA solution was microinjected into C57BL/6N embryos at the pronuclear stage, and resulting founder (F_0_) mice were analyzed by Surveyor assays as described above using tail genomic DNA. Selected founders were further characterized by DNA sequencing of the mutant allele, and mice harboring a 17-base pair deletion in exon 3 of *Arhgef3* were mated to WT mice to obtain heterozygous F_1_ progeny. Genotyping of *Arhgef3* alleles was performed as described in Figure 1A. All mouse lines were maintained on a C57BL/6N background in our institution’s animal facility.

### Muscle injury, drug injections, and animal treatments

Muscle injury was induced by injecting 50 µL of 1.2% (w/v) BaCl_2_ dissolved in saline into TA muscle. As a control, the same volume of saline was injected into the contralateral TA muscle. For systemic drug administration, triciribine (in DMSO) and chloroquine (in deionized water) stock solutions were diluted in PBS and injected intraperitoneally into mice at 1 mg/kg and 50 mg/kg, respectively. Control mice were injected with an equivalent amount of vehicle diluted in PBS. These injections were repeated every 24 hours after injury unless otherwise indicated. Lean and fat body mass were measured using an EchoMRI-700 Body Composition Analyzer (EchoMRI). All mice were housed in a room maintained at 25°C with a 12-hour light/dark cycle and received a pellet diet and water ad libitum. Mice were anesthetized with isoflurane during all surgical procedures and euthanized by cervical dislocation at the end of the experiments.

### Skeletal muscle transfection

*In vivo* transfection of TA muscle was performed by electroporation as previously described (You et al., 2018) with slight modifications. Briefly, a small incision was made on the skin covering the distal area of the TA muscle. A 30 µL of plasmid DNA solution containing either 40 µg of pCMV-Myc-ARHGEF3 or pCMV-Myc-ARHGEF3-L269E (Khanna et al., 2013) was injected into the distal end of the TA muscle with a 27-gauge needle. After the injection, two stainless steel pin electrodes (1-cm gap) connected to an ECM 830 electroporation unit (BTX/Harvard Apparatus) were laid on top of the muscle, and eight 20-ms square-wave electric pulses were delivered at a frequency of 1 Hz with a field strength of 50 V/cm. After electroporation, the incised skin was closed with a 3-0 polysorb suture.

### In situ muscle force analyses

In situ force measurement of TA muscle was performed using a 1300A Whole-Animal System (Aurora Scientific). The anesthetized mouse was placed on an isothermal stage set at 38°C, and the skin covering TA muscle and patella was incised. The distal tendon of TA muscle was tied with a 3-0 suture line and cut to isolate the muscle from tibia. After stabilizing the hindlimb by inserting a needle through a fixed post and patella tendon, the suture line was hooked onto the lever arm of the force transducer. Two electrodes were then placed on either side of the TA muscle and electrical stimulations were elicited with 0.2-ms square-wave pulses at 0.2 mA. Once muscle length was adjusted to optimal muscle length (Lo) where maximal twitch force was produced, maximum isometric tetanic force was determined in the frequency range of 50-200 Hz with 300 ms pulse duration. All tetanic contractions were separated by a 1-minute rest. Throughout the experiments, the exposed TA muscle was kept moist with a warm PBS-soaked KimWipe. Physiological cross-sectional area (pCSA) was calculated by dividing muscle mass by the product of fiber length and muscle density (1.06 g/cm^3^). Specific isometric tetanic force was calculated by dividing maximal isometric tetanic force by pCSA.

### Histochemical analyses

Isolated TA muscles were submerged in Tissue-Tek OCT compound (Sakura Finetek) at resting length and frozen in liquid nitrogen–chilled isopentane. Mid-belly cross-sections of 10 µm thickness were made with a Microm HM550 cryostat (Thermo Fisher Scientific) at −25°C, placed on microscope slides, and subjected to H&E staining. H&E-stained images randomly chosen were captured with a 20X dry objective (Fluotar, numerical aperture 0.4; Leica) on a Leica DMI 4000B microscope and analyzed for CSA of all centrally nucleated regenerating myofibers using image J (NIH). Investigators were blinded to the sample identification during all procedures.

### Western blotting

Isolated muscles were immediately frozen in liquid nitrogen and homogenized with a Polytron in ice-cold buffer A containing 50 mM Tris (pH 7.4), 0.5% sodium deoxycholate, 0.1% SDS, 1% Triton X-100, 500 mM NaCl, 10 mM MgCl_2_, 0.1 mM Na_3_VO_4_, 25 mM NaF, 25 mM β-glycerolphosphate, and 1× protease inhibitor cocktail (P8340, Sigma-Aldrich). The homogenates were pre-cleared by centrifugation at 16,200 g for 10 min (4°C). For the detection of F4/80, the frozen muscles were homogenized in ice-cold buffer B containing 20 mM Tris (pH 7.4), 0.3% Triton X-100, 2 mM EGTA, 2 mM EDTA, 0.1 mM Na_3_VO_4_, 25 mM NaF, 25 mM β-glycerolphosphate, and 1× protease inhibitor cocktail, and the homogenates were centrifuged at 800 g for 10 min (4°C) to remove myofibrillar proteins. The protein concentration in each supernatant sample was determined with the DC protein assay Kit (Bio-Rad).

For Western blotting, an equal amount of protein from each sample was boiled in Laemmli buffer, resolved on SDS-PAGE, transferred onto PVDF membrane (EMD Millipore), blocked with 5% milk in PBS-T (PBS with 0.5% Tween 20), and probed with the indicated primary and secondary antibodies according to the manufacturers’ recommendations. After washing in PBS-T, blots were developed and visualized using the SuperSignal West Pico PLUS Chemiluminescent Substrate and an iBright CL1000 Imaging System, respectively (both from Thermo Fisher Scientific). ImageJ was used to quantify each blot. The antibodies used in this study are as follows: anti-ARHGEF3 was previously reported (Khanna et al., 2013); anti-pSer473-Akt (#9271), pThr308-Akt (#9275), Akt (#9272), pThr346-NDRG1 (#3217), NDRG1 (#9408), LC3A/B (#4108), p62 (#23214), RhoA (#2117), and GAPDH (#2118) from Cell Signaling Technology; anti-pSer657-PKCα (06-822) from EMD Millipore; anti-PKCα (P4334) from Sigma-Aldrich; anti-F4/80 (clone CI:A3-1) from Bio-Rad; peroxidase-conjugated anti-rabbit (115-036-003), anti-mouse (111-036-003), and anti-rat (172-035-153) IgG antibodies from Jackson Immuno Research Laboratories.

### RhoA GEF activity assay

Muscle homogenates were prepared in buffer A as described above and an equal amount of protein from each sample was incubated with 20 µL of Rhotekin-RBD beads (Cytoskeleton) for 2 hours at 4°C and washed 4 times in ice-cold buffer containing 50 mM Tris (pH 7.4), 1% Triton X-100, 150 mM NaCl, 10 mM MgCl_2_, and 1× protease inhibitor cocktail. The beads were boiled in Laemmli buffer and active form of GTP-bound RhoA was detected by Western blotting with anti-RhoA primary antibody. RhoA-specific GEF activity was then determined by normalizing the amount of GFP-bound RhoA to the amount of total RhoA.

### Quantitative PCR (qPCR)

Frozen muscles were homogenized with a Teflon-glass homogenizer in ice-cold TRIzol (Invitrogen), and total RNA was extracted using the RNeasy Mini Kit (Qiagen) according to the manufacturer’s instructions. The purity and integrity of RNA samples were confirmed by the ratio of *A*_260_/*A*_280_ absorbance and of 28*S*/18*S* ribosomal RNA, respectively. cDNA was synthesized from 1 µg of RNA using the qScript cDNA Synthesis Kit (Quanta Bioscience) and subjected to qPCR on a StepOnePlus Real-Time PCR System (Applied Biosystems) using SYBR green. The primers used: 5’-CTTCAGCAACGAGAGAGT-3’ (forward) and 5’-GAAGGTGTCGTTGGCTTGTA-3’ (reverse) for *Arhgef3*; 5’-CCCAGTGTCTTGGCATTCTT-3’ (forward) and 5’-AGGGAAAGCAGAGGAAGCTC-3’ (reverse) for *p62*; 5’-AATCCCATCACCATCTTCCA-3’ (forward) and 5’-TGGACTCCACGACGTACTCA-3’ (reverse) for *Gapdh* as an internal control.

### Statistics

All values were presented as mean ± SEM, with individual data points shown in graphs. Sample size for each experiment was determined on the basis of previous publications and preliminary data. Mice from the same pedigree were randomly allocated to the various experimental groups except when they were used for some parameters that required a similar body weight across the groups such as CSA. Mice that showed any sign of abnormality according to pre-established criteria were excluded from experiments. A quantified sample value that deviated more than three times from the mean in a given group was considered as an outlier. Statistical significance (*P* < 0.05) was determined by two-tailed paired (when comparing to contralateral controls) or unpaired t-tests for single comparisons or one- or two-way ANOVA followed by Student-Newman-Keuls post hoc test for multiple comparisons. All statistical analyses were performed using Excel or SigmaPlot.

### Study approval

All animal experiments were performed in accordance with protocols approved by the Institutional Animal Care and Use Committee at the University of Illinois at Urbana-Champaign (#16157).

