## Supplemental Figures 1-2 for "ARHGEF3 regulates skeletal muscle regeneration and strength through autophagy"

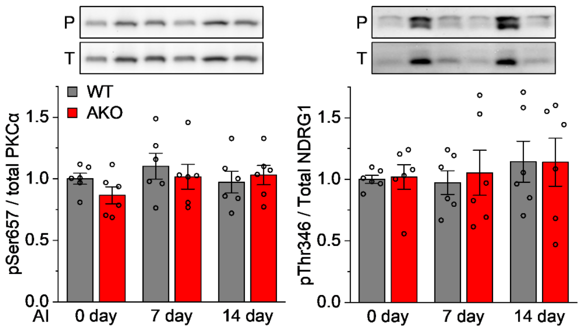


**Figure S1. Effects of ARHGEF3 KO on the phosphorylation of mTORC2 substrates. Related to Figure 3.** TA muscles from 3-month-old WT and ARHGEF3 KO (AKO) male mice were injected with $BaCl_2$ (injury) or saline (uninjured control, 0 day), collected 7 and 14 days after injury (AI), and analyzed by Western blotting for phosphorylated (p)/total protein ratio for PKCα and NDRG1 (*n* = 6).

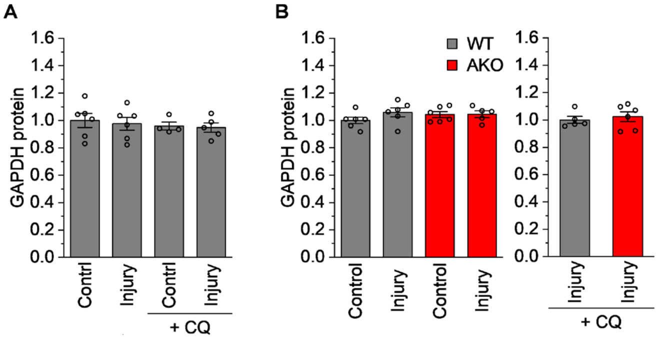


**Figure S2. Quantification of GAPDH controls. Related to Figure 5.** (**A**) Quantification of GAPDH presented in Figure 5A (*n* = 5-6). (**B**) Quantification of GAPDH presented in Figure 5B (*n* = 5-6).
